# Effect of Total Sphingomyelin Synthase Activity on Low Density Lipoprotein Catabolism in Mice

**DOI:** 10.1101/2023.02.03.527088

**Authors:** Zhiqiang Li, Mulin He, Guangzhi Chen, Tilla S. Worgall, Xian-Cheng Jiang

**Author notes:** These authors contributed equally to this work (Zhiqiang Li, Mulin He). Correspondence to Xian-Cheng Jiang, PhD, SUNY, Downstate Medical Center, 450 Clarkson Ave Box 5, Brooklyn, NY 11203.

## Abstract

**Background:** Sphingomyelin (SM) and cholesterol are two key lipid partners on cell membranes and on lipoproteins. Many studies have indicated the influence of cholesterol on SM metabolism. This study examined the influence of SM biosynthesis on cholesterol metabolism.

**Methods:** Inducible global *Sms1* KO/global *Sms2* KO mice were prepared to evaluate the effect of whole-body SM biosynthesis deficiency on lipoprotein metabolism. Tissue cholesterol, SM, ceramide, and glucosylceramide levels were measured. TG production rate and LDL catabolism were measured. Lipid rafts were isolated and LDL receptor mass and function were evaluated. Also, the effects of exogenous sphingolipids on hepatocytes were investigated.

**Results:** We found that total SMS depletion significantly reduced plasma SM levels. Also, the total deficiency significantly induced plasma cholesterol, apoB, and apoE levels. Importantly, total SMS deficiency, but not SMS2 deficiency, dramatically decreased LDL receptors in the liver and attenuated LDL uptake through the receptor. Further, we found that total SMS deficiency greatly reduced LDL receptors in the lipid rafts which contained significantly lower SM and significantly higher glucosylceramide as well as cholesterol. Furthermore, we treated primary hepatocytes and Huh7 cells (a human hepatoma cell line) with SM, ceramide, or glucosylceramide, and we found that only SM could up-regulate LDL receptor levels in a dose-dependent fashion.

**Conclusions:** Whole-body SM biosynthesis plays an important role in LDL-cholesterol catabolism. The total SMS deficiency, but not SMS2 deficiency, reduces LDL uptake and causes LDL-cholesterol accumulation in the circulation. Given the fact that serum SM level is a risk factor for cardiovascular diseases, inhibiting SMS2 but not SMS1 should be the desirable approach.

**Graphic Abstract:** 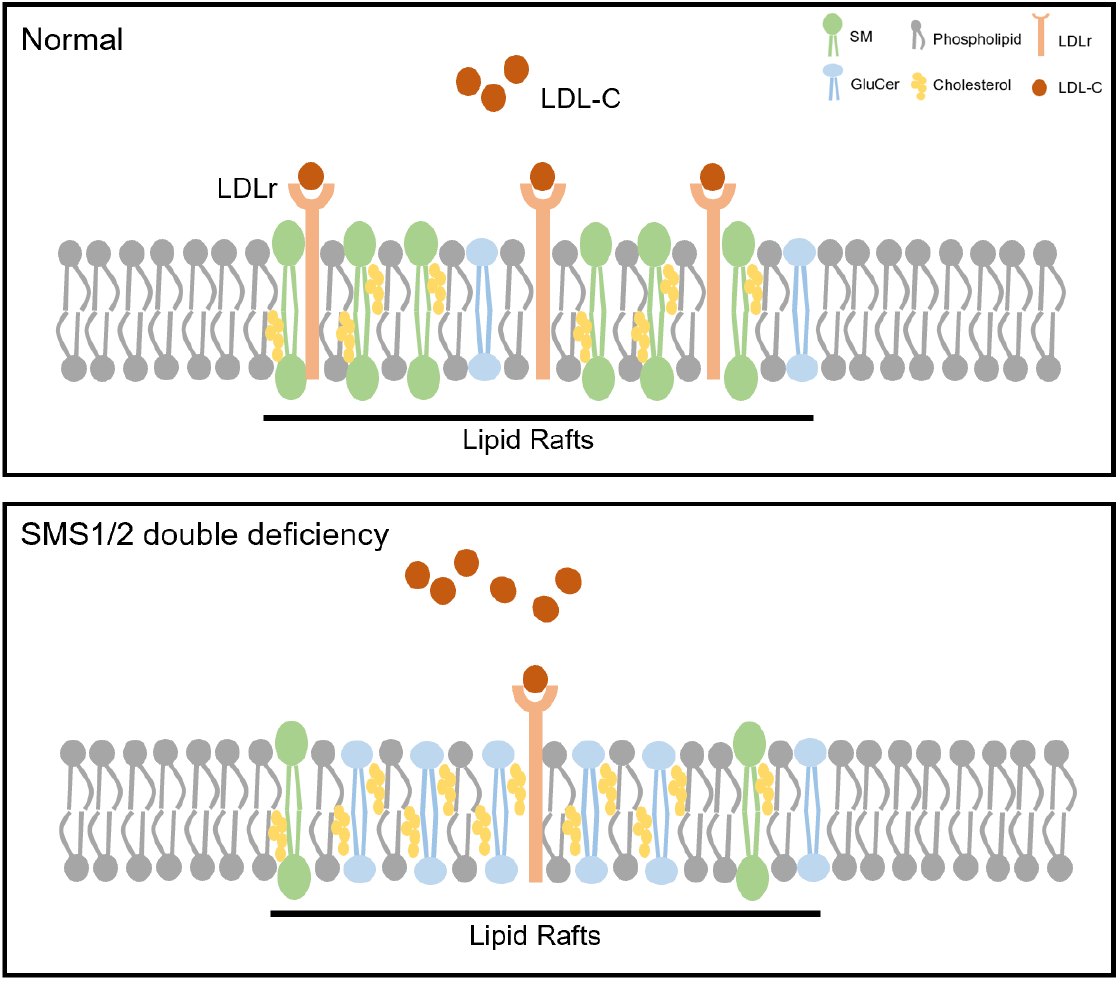

## Introduction

Sphingolipid biosynthesis in the ER starts with the condensation of serine and palmitoyl-CoA to form 3-ketosphinganine, catalyzed by serine palmitoyltransferase (SPT), which is the key enzyme for sphingolipid biosynthesis. Perturbations of SPT activity have been linked to metabolic diseases. Park et al.^1^ we^2^ and Glaros et al.^3^ reported that treatment with myriocin, an inhibitor of SPT activity, decreases atherosclerosis induced by a high-fat/high-cholesterol diet in mouse models. The mechanism could involve the reduction of plasma sphingomyelin (SM), ceramide, and glycosphingolipids. In mice, myriocin-mediated reduction of ceramide protects against diet-induced insulin resistance ^4^, and dyslipidemia ^5^. Although myriocin can help prevent metabolic diseases, it is toxic and has unfavorable physicochemical properties such as low solubility^6^. Thus, we need other approaches to regulate sphingolipid synthesis.

There are three steps after SPT to form ceramide, a hub molecule for the biosynthesis of all sphingolipids, including sphingomyelin (SM)^7^. Human serum SM level is an independent risk factor for cardiovascular disease^8^. SM is accumulated in human and animal atheroma^9^. LDL in human atheroma is enriched with SM^10^. Plasma SM level in *Apoe* KO mice is 4-fold higher than that in wild-type (WT) mice^11^, and this contributes to the development of atherosclerosis^12^. SM increases from 10% at birth to 60% in patients who had atherosclerotic plaques, and SM contributes to sudden death^13^. Also, SM level is one of the factors associated with an increased risk of myocardial infarction^14^. SM is one of the important lipids on the monolayer of plasma lipoproteins^15^, including LDL, SM levels should have an important impact on LDL metabolism. Thus, manipulation of SM biosynthesis could provide a clue for this.

SM synthase (SMS) family has three members, they are SMS1, SMS2, and SMSr (SMS related protein). Both SMS1 and SMS2 but not SMSr catalyze the conversion of ceramide to SM^16^. SMS1 and SMS2 are co-expressed in all tested tissues, including the liver, small intestine, adipose tissue, muscle, and so on. Thus, neither SMS1 nor SMS2 deficient approach is sufficient to evaluate the effect of SM reduction *in vivo*. Previously, we found that global *Sms1* knockout (KO) mice, with a noticeable problem of postnatal lethality, have significantly less SM in the plasma, liver, and macrophages but have only a marginal effect on ceramide^17^. We also prepared global *Sms2* KO mice and found the mice are healthy and have reduced SM in the circulation^18^. Moreover, we prepared albumin promoter driven liver-specific *Sms1* KO/global *Sms2* KO mice to evaluate the effect of liver SM biosynthesis in lipoprotein metabolism. Our results showed that liver total SMS blocking can reduce VLDL production^19^. Since SMS1 is depleted in the early stage of life, some compensations occurred in response to SMS1 deficiency at that time. Thus, previous approaches, using either global or liver-specific *Sms1* KO mice, may not reflect the situation in adulthood and may not provide useful information for disease treatment. Thus, in the current study, we evaluated lipoprotein metabolism in inducible global SMS-deficient mice and we found that whole-body SMS deficiency has an important impact on LDL catabolism.

## Materials and Methods

The data that support the findings of this study are available from the corresponding author upon reasonable request.

### Animals

In this study, all animal experiments were conducted with the approval of the Institutional Animal Care and Use Committee (IACUC) at SUNY Downstate Health Sciences University. Both male and female mice were used. Wild-type (WT) C57BL/6J mice were obtained from The Jackson Laboratory (Strain #:000664). The strategy to prepare SMS2^−/−^ SMS1^fl/fl^ (sKO) mice was previously described^27^. The SMS2^−/−^ SMS1^fl/fl^ UBC-Cre-ER^T2^ (dKO) mice were generated by crossing SMS2^−/−^ SMS1^fl/fl^ (sKO) mice with UBC-Cre-ER^T2^ transgenic mice from The Jackson Laboratory (Strain #: 008085) described as **Supplemental Figure 1**. Tamoxifen (80 mg/kg body weight, dissolved in corn oil) was intraperitoneally injected into mice at the age of 8-12 weeks to induce the ubiquitin C (UBC) promoter mediated Cre recombination. Five alternative days of tamoxifen injection generated global inducible SMS1 and SMS2-knockout mice. WT and sKO mice were also injected with an equal volume of tamoxifen. Mice fed with anormal laboratory diet (PicoLab Rodent Diet 5053; protein 20.0%, fat 4.5%, fiber 6.0%) were used for all experiments. Overnight fasted mice were sacrificed on Day 17 after tamoxifen injection, and organs were collected and immediately frozen in liquid nitrogen and stored at −80°C. All mice have C57BL/6J genetic backgrounds.

### Quantitative reverse transcription PCR

Tissues were homogenized, and total RNA was isolated using TRIzol (Invitrogen) according to the manufacturer’s instructions. Two micrograms of total RNA were used to produce cDNA using high-capacity cDNA reverse transcription kits (Applied Biosystems). Real-time PCR was performed using the SYBR Green Master Mix System (Applied Biosystems) on the StepOnePlus Real-time PCR system (Applied Biosystems). PCR amplification was performed in triplicated on 96-well optical reaction plates and replicated at least three times. The ΔΔCt method was used to analyze the relative changes in gene expression from qRT-PCR. The Ct of mouse GAPDH or 36B4 cDNA were used to normalize all samples. Primers are listed in Major Resource Table.

### SMS activity assay

The SMS activity assay procedure was similar as described before^20^. Briefly, mouse liver (0.2 g) was homogenized in 50 mM Tris-HCl (pH 7.4), 1 mM EDTA, and 250 mM sucrose. For *SMS* activity, each reaction (total volume 600 μl) contained 50 mM Tris-HCl (pH 7.4), 25 mM KCl and 1% fatty acid-free BSA, 3 μl phosphatidylcholine (Sigma, P6638) (stock solution 20 mg/ml), 2 μl NBD-Ceramide (Sigma, N8278) (stock solution 1 mg/ml), and 50 μg of protein. The reaction was incubated at 37 °C for 1.5 hours and the reactions were terminated by adding chloroform/methanol (2:1, v/v) vigorously. The lower organic phase was collected after spin at 6000 × *g* for 10 min and dried under nitrogen gas. Lipids were dissolved in 20 μl chloroform and then were applied to a TLC plate. The running solvent was chloroform/ methanol/ 20% NH4OH (14:6:1, v/v/v). The fluorescence signal was detected under UV and the intensity was quantified by NIH Image software.

### Plasma and liver lipid measurements

Mice were fasted overnight before plasma and tissue collection. Cholesterol, phospholipids, TG, free cholesterol, and non-esterified fatty acids (NEFA) were measured using commercially available kits following the manufacturer’s protocols (Wako Pure Chemical Corporation; Thermo Scientific). To minimize technical variances, we randomly chose 4 male or female individual mice from each group and collected their fasted plasma samples to run fast protein liquid chromatography (FPLC) in turn. 200 μl of pooled plasma samples were chromatographed by using a Superose 6 Increase 10/30 GL Column (29091596, Cytiva).

### Sphingolipids analyses by liquid chromatography with tandem mass spectrometry (LC-MS/MS)

Sphingolipid levels were quantified in mice plasma, liver homogenates, and lipid rafts fractions by targeted high pressure liquid chromatography electrospray ionization tandem mass spectrometry (HPLC-MS/MS) in positive multiple reaction monitoring (MRM) mode using minor modification of a described method as we performed before^21,22^. Briefly, the method was validated for 5 dihydroceramides: (d18:0/16:0 d18:0/18:0, d18:0/18:1, d18:0/24:0, d18:0/24:1), 6 ceramides (d18:1/C16:0, d18:1/C18:0, d18:1/C20:0, d18:1/C22:0, d18:1/C24:0, d18:1/C24:1), 4 sphingomyelins (SM d18:1/C16:0, SM d18:1/C18:0, SM d18:1/C18:1, SM d18:1/C24:1), and 4 long-chain bases: sphingosine (SO d18:1), sphinganine (SA d18:0), sphingosine-1-phosphate (S1P d18:1), sphinganine-1-phosphate (Sa-1-P d18:0). 25 μl plasma, tissue or cell extracts (50-200 μg protein depending on sphingolipid content) were extracted by vortexing 30 min in 500-1000 μl dichloromethane / methanol (1:1) with addition of 25 pmol each of internal standards (N-lauroyl-D-erythro-sphingosine and N-lauroyl-D-erythro-sphingosyl phosphoethanolamine (C17 base)). Samples were centrifuged (4000 g, 10 min) to precipitate cell debris. An aliquot of 300 μl was transferred into a new organic solvent resistant (2 ml × 96-well) plate from which 3 μl are injected into an Agilent 1200 HPLC equipped with Agilent Poroshell 120 EC C18(2.1 × 50 mm 2.7 micron) column linked to an Agilent 6430 triple quadrupole mass spectrometer. Mobile phase A consisted of methanol/water/chloroform/formic acid (55:40:5:0.4 v/v); Mobile phase B consisted of methanol/acetonitrile/chloroform/formic acid (48:48:4:0.4 v/v). After holding mobile 95 % Mobile phase A for 0.8 min, the gradient was gradually increased to 95% mobile phase B at 1.4 minutes and held up to 7.2 min. The column was then equilibrated with mobile phase A from 7.2 to 7.5 min. With a flow rate was 0.6 mL/min, the duration of the entire run was 9.65 min. Mass Hunter optimizer and pure synthetic standards (Avanti Polar Lipids) were used to determine optimum fragmentation voltage, precursor/ product ions and m/z values. Peak calls and abundance calculations were obtained with MassHunter Workstation Software Version B.06.00 SP01/Build 6.0.388.1 (Agilent). Final concentrations were calculated from a standard curve for each sphingolipid run in parallel.

### Hepatic TG production

Female mice (n=5-7 per group) were fasted overnight and then injected with poloxamer 407 intraperitoneally (1 g/kg BW) (Sigma, 16758). Plasma TG concentrations were measured at 0, 1, and 2 h following injection. For 1h and 2 h TG measurements, plasma was diluted 20 times with PBS.

### Immunoblotting

Tissue homogenates or cell lysates were prepared in RIPA lysis buffer supplemented with protease inhibitors (Roche) and subjected to Western blotting as previously described^30^. The following primary antibodies were used: ApoB (Abcam, 20737); ApoE (Santa Cruz, sc-6384); ApoA1 (Proteintech, 14427-1-AP); Pre-albumin (Proteintech, 11891-1-AP); LDLr (Proteintech, 10785-1-AP); GAPDH (Novus Biologicals, 300-324); SR-B1 (Novus Biologicals, 400-101); ABCA1 (Novus Biologicals, 400-105); ABCG5 (Proteintech, 27722-1-AP); BSEP (Invitrogen, PA5-78690); PCSK9 (Proteintech, 27882-1-AP); Lyn (Santa Cruz, sc-15); Caveolin-1 (Cell Signaling, 3238); β-actin (Abcam, 8227). The blotting membrane was also stained with Ponceau S (Sigma, P7767) for total protein normalization.

### Lipid rafts isolation

The procedure of liver lipid raft extraction was similar as previously described^28^. Briefly, mouse liver was homogenized in 0.5 M Na_2_CO_3_ buffer (pH 11) containing protease inhibitor, and then centrifuged at 1,300 g for 5 minutes to pellet cellular debris and nuclei. One milliliter of the postnuclear supernatant was mixed with 1 ml of 90% sucrose solution in the buffer containing 50mM HEPES (pH 6.5), and 150 mM NaCl, placed at the bottom of the ultracentrifuge tube. A discontinuous sucrose gradient was formed by overlaying with 6 ml of 35% sucrose and 3.5 ml of 5% sucrose in the same buffer containing 0.25 M Na_2_CO_3_ and centrifuged at 38,000 rpm for 18 h in an SW41 rotor (Beckman). Twelve fractions were collected from top to bottom. The fractions were dissolved in 5x SDS loading buffer for immunoblotting analysis.

### LDL uptake and LDL receptor immunostaining assay

LDL uptake and cellular LDL receptor expression were determined according to the manufacturer’s protocol (Abcam, 133127). Briefly, primary hepatocytes were isolated from 8-week-old WT, sKO, and dKO mice (male and female), plated in 8-well glass chamber slides (Nunc Lab-Tek, 154534), and cultured in DMEM medium supplemented with 1% BSA (essentially fatty acid free). After 24hrs incubation, the culture medium was replaced with 75 μl/well LDL-Dylight 550 solution, followed by 4 hrs incubation at 37°C in a 5% CO2 incubator, and LDL uptake was analyzed with fluorescent microscope. Subsequently cells were fixed for LDL receptor immunofluorescent staining using anti-LDL receptor antibody and Dylight 488 secondary antibody. Cells were counterstained for nuclear DNA with DAPI (1:1000). Rabbit mAb IgG isotype control (Cell signaling, 3900) was used as a negative control. To minimize technical variations, all samples were stained and imaged at the same time. To minimize observer error, images were chosen and analyzed in a blinder manner. Immunofluorescent images were captured using an Olympus BX53 immunofluorescent microscope. Mean fluorescence intensity of LDLr staining was determined using Image J software with n=5 or more and normalized to the total number of DAPI stained nuclei in the fields of view.

### Exogenous Ly93, sphingomyelin, ceramide, and glucosylceramide supplementation

Ly93, aSMS inhibitor (a gift from Dr. Deyong Ye, School of Pharmacy, Fudan University), was dissolved in DMSO. SM mixture including 24:1, 24:0, and 16:0 (1:1:1) (Avanti Polar Lipids, Inc) was dissolved in ethanol. Ceramide mixture including 24:1, 24:0, and 22:0 (1:1:1) (Avanti Polar Lipids, Inc) was dissolved in ethanol/dodecane (98:2, v/v). Glucosylceramide (Avanti Polar Lipids, Inc) stock solution was prepared as described^31^. Human liver hepatoma 7 (Huh7) cells or primary hepatocytes derived from *Sms2* KO (sKO) mice (male or female) were plated in collagen-coated 6-well plates and incubated with different concentrations of exogenous of Ly93 for 18 hrs, or SM mixture, ceramide mixture, glucosylceramide, or vehicle for 28 hours. We have not seen significant sex differences in the results with isolated primary hepatocytes from mice (detailed sex information of the used mice were interpretated in each figure legend). In all experiments, cells were cultured with DMEM medium containing 1% essentially fatty acid free BSA and 0.1% penicillin-streptomycin. Cell lysates and extracted plasma membranes were prepared for immunoblotting analysis.

### Statistics

Data are presented as mean ± SEM. Each experiment was repeated at least three times. The sample number in each experimental group and mice sex information are noted in the figure legends. All data were assessed for normality and equal variance to determine parametric or nonparametric test. For comparison of two groups, data that did not pass the test for normality and equal variance were analyzed by the Mann-Whitney rank sum test. Weight analysis were analyzed by repeated-measures 2-way ANOVA with Tukey multiple comparisons test. Data from multiple groups were analyzed by 1-way ANOVA followed by Sidak multiple comparisons test, Kruskal-Wallis test followed with Dunn’s multiple comparisons test or 2-way ANOVA with Tukey multiple comparisons test to evaluate the significance for comparisons with 1 or 2 independent variables using GraphPad Prism software. Data with *p*<0.05 were considered statistically significant.

## Results

### Whole-body depletion of *Sms1* and *Sms2*

To investigate both SMS1 and SMS2 functions, we generated inducible whole-body *Sms1* and *Sms2* double knockout (KO) mice (SMS2^−/−^ SMS1^fl/fl^ UBC-Cre-ER^T2^) (**Supplemental Figure 1A**). Control and *Sms2* KO littermate mice received an equal volume of tamoxifen for five alternative days (**Figure 1A**). At day 17, we measured *Sms1* mRNA levels in female mice. We found that the mRNA reduction in the double KO (dKO) mice were achieved in over 90% of all tested tissues, including liver, small intestine, adipose tissue, lung, and muscle (**Figure 1B**) and that liver SM synthase activity was completely blocked (**Figures 1C and 1D**). Moreover, we found that the dKO mice exhibited a significant reduction in body weight as early as day 8 following tamoxifen injection compared to control and *Sms2* KO mice, and the reduction became more significant over the course of the study (**Figure 1E**). We also examined *Sms2* mRNA levels in different tissues and they have no expression at all (**Supplemental Figure 1B**). Male mice have similar phenomena (data not shown).

**Figure 1.**
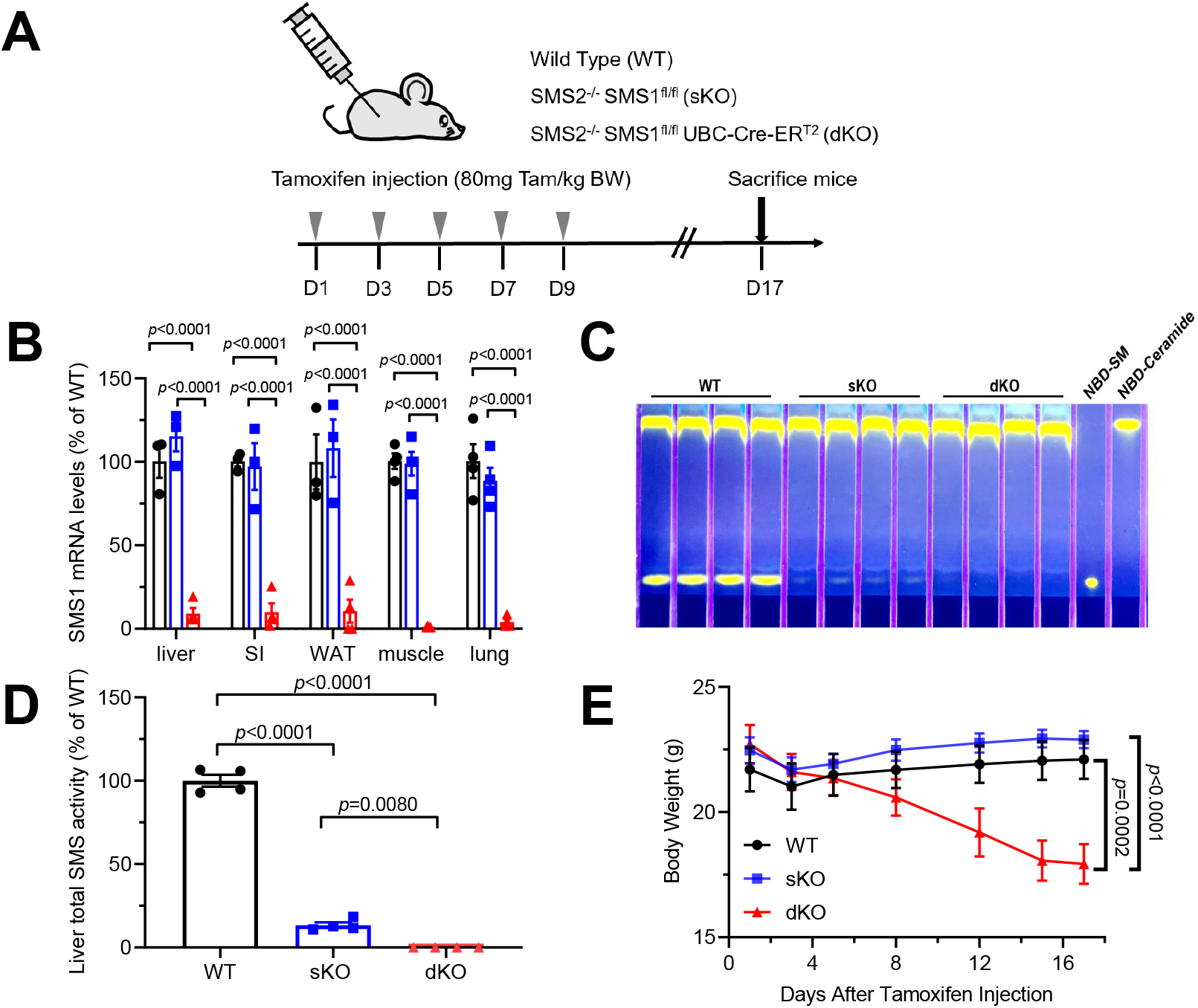
Whole-body depletion of *Sms1* and *Sms2*. Panel A, Scheme of tamoxifen (Tam) injection. Panel B, *Sms1* gene expression in liver, small intestine, white adipose tissue, muscle, and lung from wild-type (WT), *SMS2*^−/−^(sKO), and *SMS2*^−/−^/*SMS1^fl/fl^ UBC-Cre-ER^T2^* (dKO) mice 17 days after the first tamoxifen injection (female, n=4 individual mice per group). Statistical analysis was done using 2-way ANOVA followed by Tukey multiple comparisons test. Panel C, Liver total SMS activity (female, n=4 individual mice per group). Panel D, Quantification of liver total SMS activity. Statistical analysis was done using 1-way ANOVA followed by Sidak multiple comparisons test. Panel E, Body weight of mice fed a normal laboratory diet (female, n=8-10 per group). Statistical analysis was done using two-way ANOVA for repeated measurements followed by Tukey post hoc test.

### Effects of inducible SMS deficiency on plasma lipids and apolipoproteins

We sought to measure plasma lipid levels in these mice. Firstly, we utilized LC/MS/MS to evaluate SM and ceramide levels. As expected, we found that all tested subspecies of SM were significantly decreased in the dKO mice (**Figure 2A**). Of interest, all subspecies of ceramide were significantly increased in *Sms2* KO mice, however, only 16:0 and 18:0 ceramides were further increased in the dKO mice (**Figure 2B**). Further, we found that plasma triglyceride levels were significantly decreased in the inducible dKO mice compared with wild type (WT) and *Sms2* KO mice (**Figures 2C**) which was same as that of liver-specific *Sms1*/global *Sms2* dKO mice^19^. We then evaluated liver triglyceride production rate using Poloxamer 407 to block lipoprotein lipase as we did before^23^. The deficiency significantly reduced triglyceride production (**Figure 2D, Supplemental Figure 2A**). These observations were similar to what we have observed in liver-specific *Sms1*/global *Sms2* dKO mice^19^. However, unexpectedly, we found that total cholesterol levels were significantly increased in the inducible dKO female and male mice in a time-dependent fashion, compared with wild type and *Sms2* KO mice (**Figure 2E, Supplemental Figure 2B**). Moreover, total choline-containing phospholipids were also significantly increased (**Figure 2F, Supplemental Figure 2C**). Given the fact that SM is one of the choline-containing phospholipids, the increase of total phospholipids indicates a dramatic increase in phosphatidylcholine.

**Figure 2.**
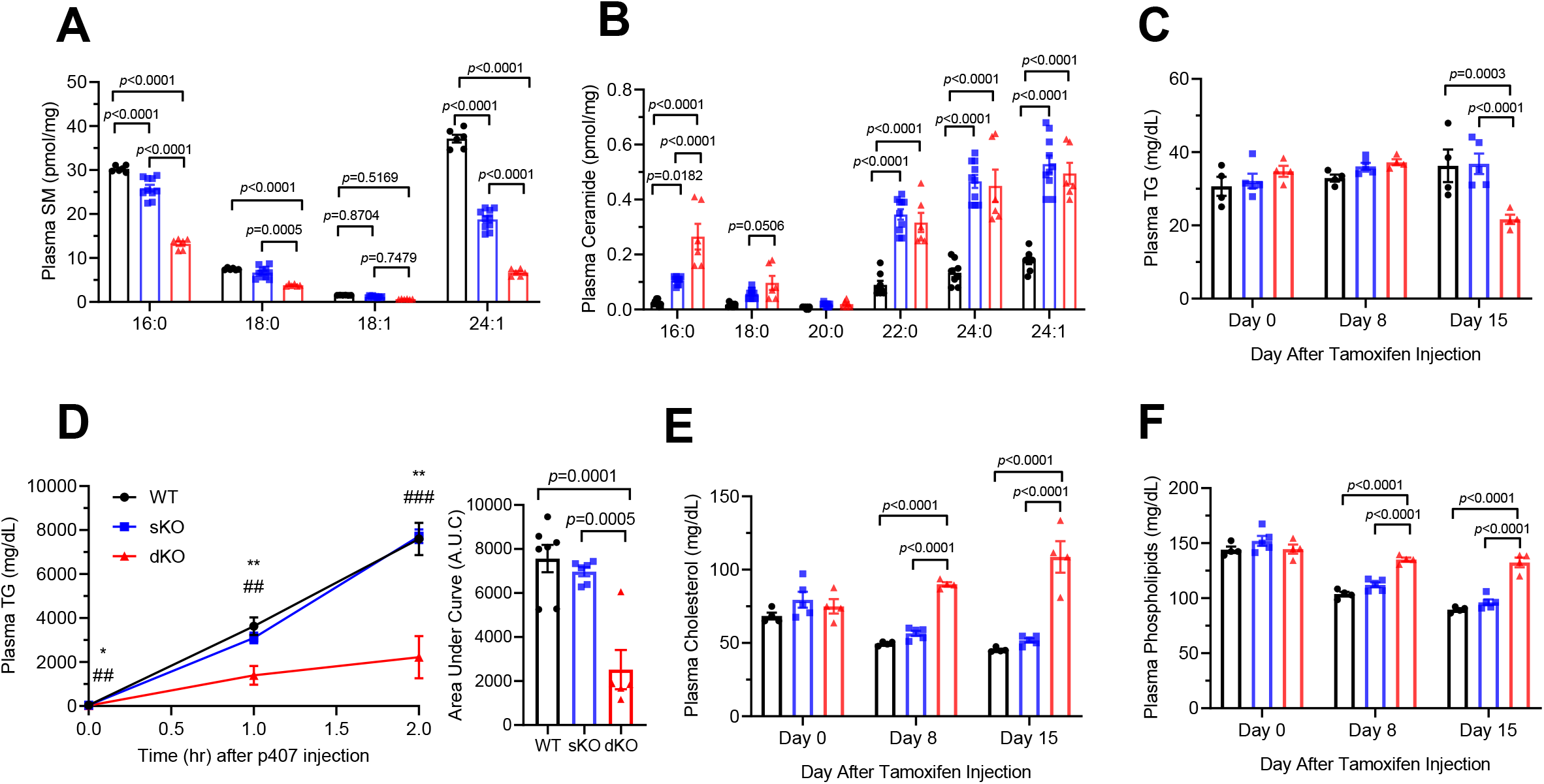
Effect of inducible *SMS* deficiency on plasma lipids. Panels A-B, Plasma sphingomyelin, and ceramide were measured by LC/MS/MS. Plasma was collected from 4h fasting wild-type (WT), *SMS2*^−/−^(sKO), and *SMS2*^−/−^/*SMS1^fl/fl^ UBC-Cre-ER^T2^* (dKO) mice on Day 17 after tamoxifen injection (female, n=6-10 individual mice per group). Panel C, Plasma triglyceride levels (female, n= 4-5 individual mice per group). Panel D, TG production rate measurement and its area under curve (A.U.C). WT, *SMS2^−/−^*, and the dKO mice were treated with poloxamer 407 (1 mg/g of body weight, i.p.) on Day 17 after tamoxifen injection, and plasma TG was measured at 0, 1, and 2h (female, n=5-7 individual mice per group). ** *p*< 0.01, WT or *SMS2^−/−^* vs the dKO. Panels E-F, Plasma cholesterol, and phospholipids profiles before and after tamoxifen injection (female, n= 4-5 individual mice per group). Panel A through C and E through F, Statistical analysis was done using 2-way ANOVA followed by Tukey multiple comparisons test. Panel D, Statistical analysis was done using 1-way ANOVA followed by Sidak multiple comparisons test.

Next, we utilized fast protein liquid chromatography (FPLC) and pooled mouse plasma to confirm the above observations. Indeed, the dKO mice had lower triglyceride levels on non-HDL fraction (**Figure 3A**) and higher cholesterol levels on both non-HDL and HDL fractions (**Figure 3B**). Then, we measured apolipoprotein levels using immunoblots. We found that the inducible dKO mice exhibited a significant increase in apoB (**Figures 3C and 3D**), apoE (**Figures 3C and 3E**) but not apoA1 (**Figures 3C and 3F**) levels compared with that of wild type and *Sms2* KO mice. Collectively, cholesterol homeostasis is disrupted when both SMS1 and SMS2 are depleted globally and simultaneously, thereby cholesterol is accumulated in the circulation.

**Figure 3.**
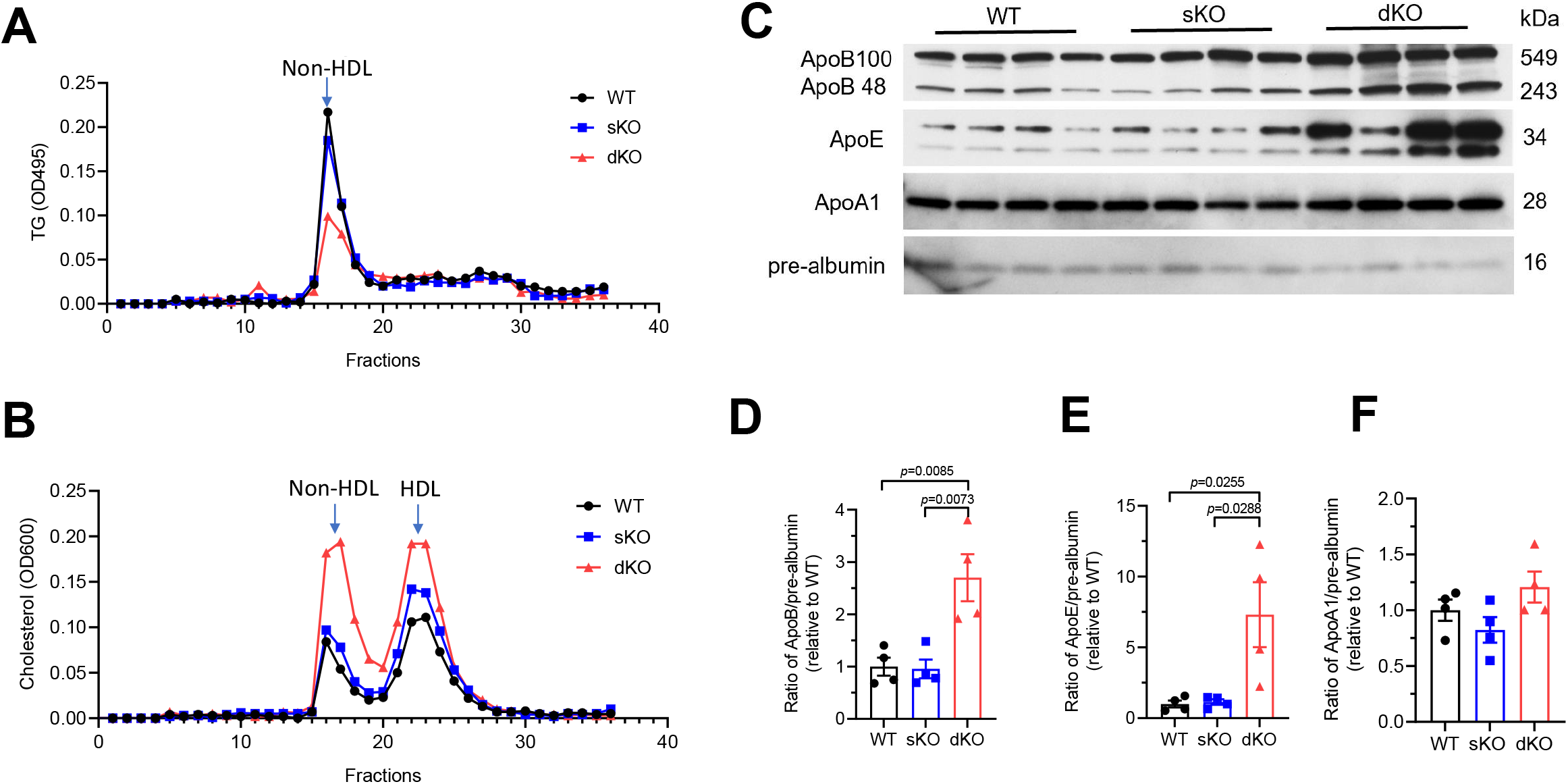
Effect of inducible SMS deficiency on plasma lipoproteins. Panels A-B, FPLC for plasma triglyceride and cholesterol distribution analysis, using pooled plasma from 4h fasting wild-type (WT), *SMS2^−/−^* (sKO), and *SMS2^−/−^/SMS1^fl/fl^ UBC-Cre-ER^T2^* (dKO) mice on Day 17 after tamoxifen injection (female, n=5-7 individual mice per group). Panel C, Western blot analysis of plasma ApoB, ApoE, and ApoA1 (female, n=4 individual mice per group). Panel D-F, Quantification of all tested apolipoproteins. Panel D through F, Statistical analysis was done using 1-way ANOVA followed by Sidak multiple comparisons test.

### Effects of inducible SMS deficiency on liver lipids

To investigate how inducible and global SMS deficiency causes cholesterol accumulation in plasma, we focused on the liver, which is considered the major site for maintaining whole-body cholesterol homeostasis. We first accessed liver sphingolipid levels and found all tested SM in liver homogenates were dramatically reduced compared to wild type and *Sms2* KO as expected (**Figure 4A**). Consistent with the finding in liver-specific *Sms1*/global *Sms2* dKO mice^24^, we observed that glucosylceramides were accumulated in the liver (**Figure 4B**), which could be directly linked with liver abnormality reflected by increasing plasma alanine aminotransferase (ALT) levels (**Figure 4C**). Surprisingly, ceramide, as a substrate of SMS activity, was reduced in the liver (**Figure 4D**), which is opposite to the plasma ceramide levels (**Figure 2B**). Interestingly, we also found that cholesterol was also accumulated in the liver of the dKO mice (**Figure 4E**) but no major differences in liver phospholipids (**Figure 4F**).

**Figure 4.**
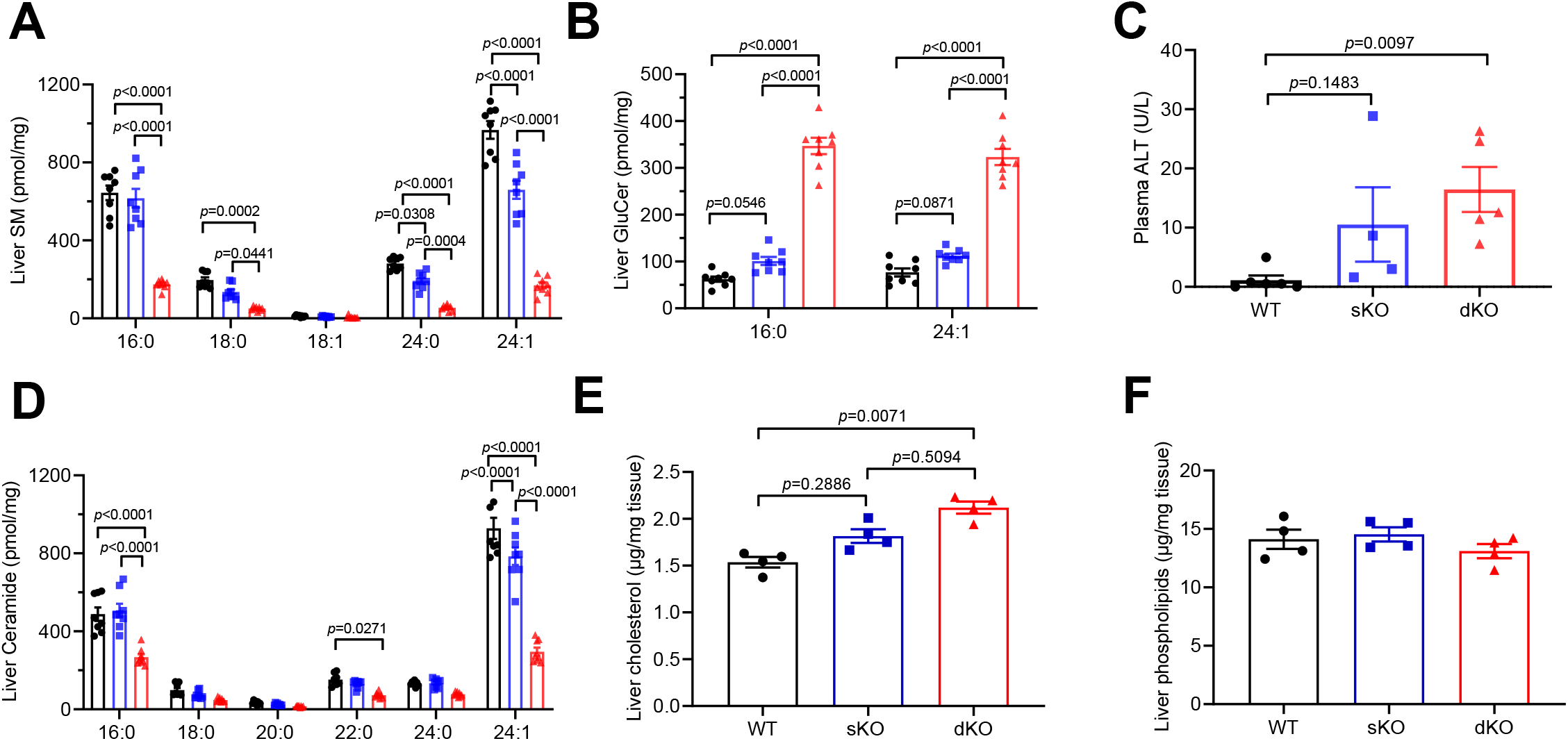
Effect of inducible SMS deficiency on liver lipids. Wild-type (WT), *SMS2^−/−^* (sKO) and *SMS2^−/−^ SMS1^fl/fl^ UBC-Cre-ER^T2^* (dKO) mice were sacrificed on Day 17 after tamoxifen injection, and livers were collected. Panels A-B, Liver sphingomyelin, and glucosylceramide levels, measured by LC/MS/MS (female, n=8 individual mice per group). Panel C, Plasma ALT levels (male, n=4-6 individual mice per group). Panel D, Liver ceramide levels measured by LC/MS/MS (female, n=8 individual mice per group). Panels E-F, Liver cholesterol, and phospholipids levels (female, n=4 individual mice per group). Panel A, B and D, Statistical analysis was done using 2-way ANOVA followed by Tukey multiple comparisons test. Panel C, E and F, Statistical analysis was done using Kruskal-Wallis test followed with Dunn’s multiple comparisons test.

### Effects of inducible SMS deficiency on liver LDL receptor expression and LDL uptake

To further characterize the effects of inducible SMS deficiency, we accessed protein levels of LDL receptor, scavenger receptor B1 (SR-BI), bile salt export pump (BSEP), ATP binding cassette AI (ABCA1), ABCG5, and PCSK9, all these proteins are involved in cholesterol homeostasis. The first interesting observation was that LDLr protein expression was reduced in the livers of the dKO mice, male and female, compared with wildtype and *Sms2* KO mice (**Figure 5A**), while no significant differences were observed in other protein levels (**Figure 5B**). We also found that SMS1 deficiency alone significantly reduced LDLr levels in the liver (**Supplemental Figure 3**). To validate our results obtained from mouse liver tissues, we isolated and cultured primary hepatocytes from wildtype, *Sms2* KO and *Sms1/Sms2* mice. We utilized fluorescence (Dylight 550) labeled LDL and anti-LDL receptor antibody to examine LDL uptake and the expression of cellular LDL receptor. As expected, LDL receptor mass, analyzed through immunofluorescence, was greatly reduced in the dKO hepatocytes (**Figures 5C and 5D, Supplemental Figure 4**). Consistently, the reduction of LDL receptor contributed to the reduction of intercellular LDL (**Figure 5C**). Collectively, these results confirmed that the accumulation of plasma cholesterol observed in the dKO mice is due to the decreased ability of LDL uptake in the liver through the downregulation of LDL receptor.

**Figure 5.**
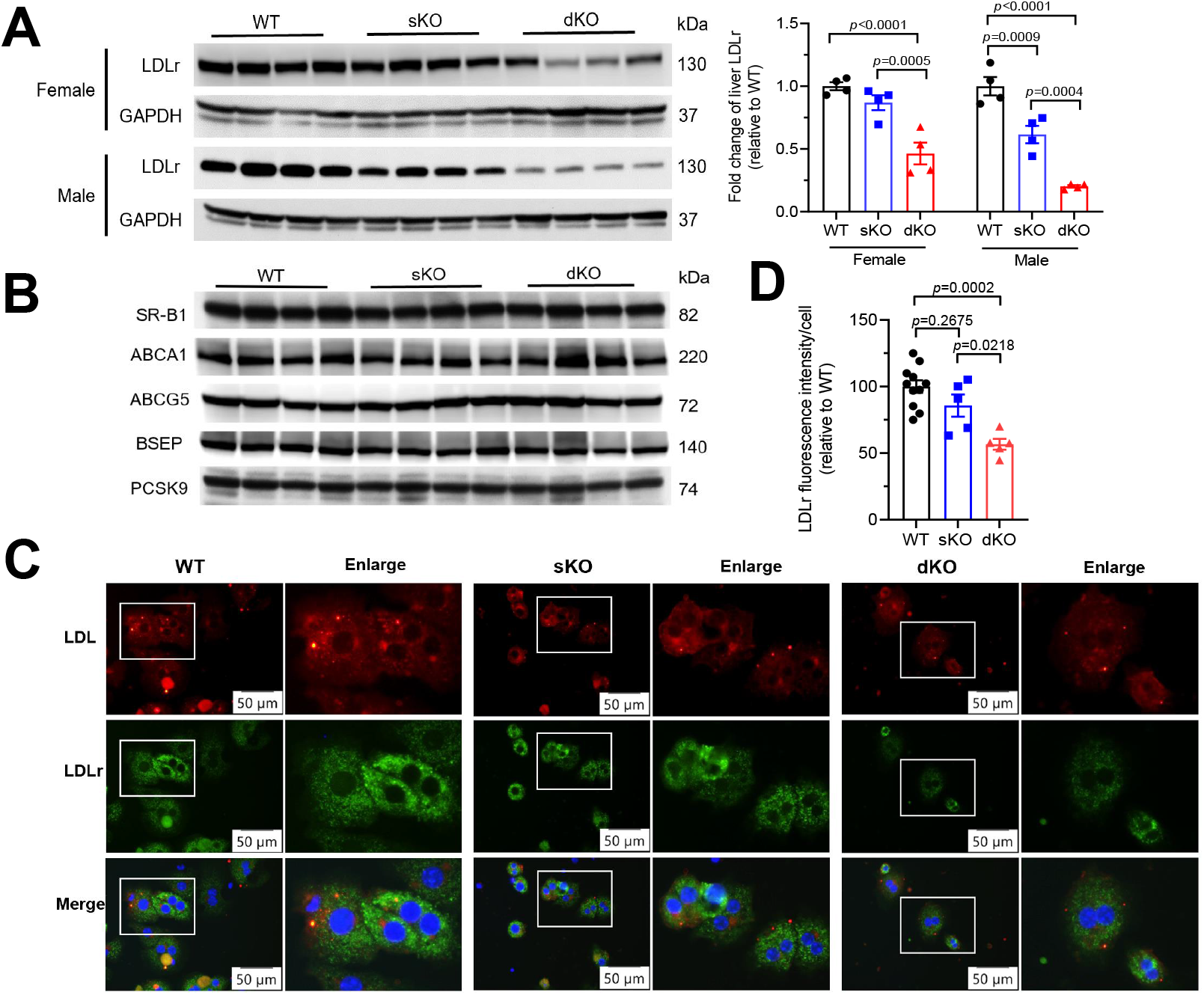
Liver LDL receptor expression and LDL uptake are decreased in inducible *Sms1/Sms2* deficient mice. Panel A, Western blot of LDL receptor expression in livers of wild-type (WT), *SMS2^−/−^ SMS1^fl/fl^* (sKO), and *SMS2^−/−^ SMS1^fl/fl^ UBC-Cre-ER^T2^* (dKO) mice that were sacrificed on Day 17 after tamoxifen injection (Female and male, n=4 individual mice per group). Panel B, Western blots for SR-B1, ABCA1, ABCG5, BSEP and PCSK9 in mice liver homogenates (female, n=4 individual mice per group). Panel C, Representative images of LDL uptake in primary hepatocytes isolated from WT, sKO, and dKO mice. Three independent experiments in triplicate. Red fluorescence denotes LDL; Green fluorescence denotes LDL receptor; Blue fluorescence denotes DAPI. Scale bars: 50 μm. Panel D, Quantification of LDL receptor fluorescence intensity/cell (n=6-10 individual mice per group). Panel A, Statistical analysis was done using 2-way ANOVA followed by Tukey multiple comparisons test. Panel D, Statistical analysis was done using 1-way ANOVA followed by Sidak multiple comparisons test.

### Reduction of SM level decreases LDL receptor in hepatocyte membrane lipid rafts

SM is one of the major lipids that contribute to the organization of lipid rafts, which are cell membrane microdomains responsible for many receptors mediated cell functions. The SMS-deficiency mediated reduction of SM in the liver could have impact on the rafts where the LDL receptor is located^25^. To validate our hypothesis, we first isolated the lipid rafts from the liver. As indicated in **Figure 6A**, the isolated lipid rafts lined very well with two markers, caveolin-1 and Lyn kinase. Two cholesterol peaks were identified in the lipid rafts (4-6) and non-rafts (10-12) fractions (**Figure 6B**). We then measured SM subspecies in the rafts and non-rafts and found that, indeed, SM was greatly enriched in the rafts. As expected, all tested SM in lipid rafts were dramatically decreased compared to wild type and *Sms2* KO (**Figure 6C**). We also measured glucosylceramide and found that it was dramatically increased in lipid rafts of dKO mouse livers (**Figure 6D**). Moreover, both rafts and non-rafts regions contained ceramide, with higher levels in the former, and SMS double deficiency caused significant reduction of it (**Figure 6E**).

**Figure 6.**
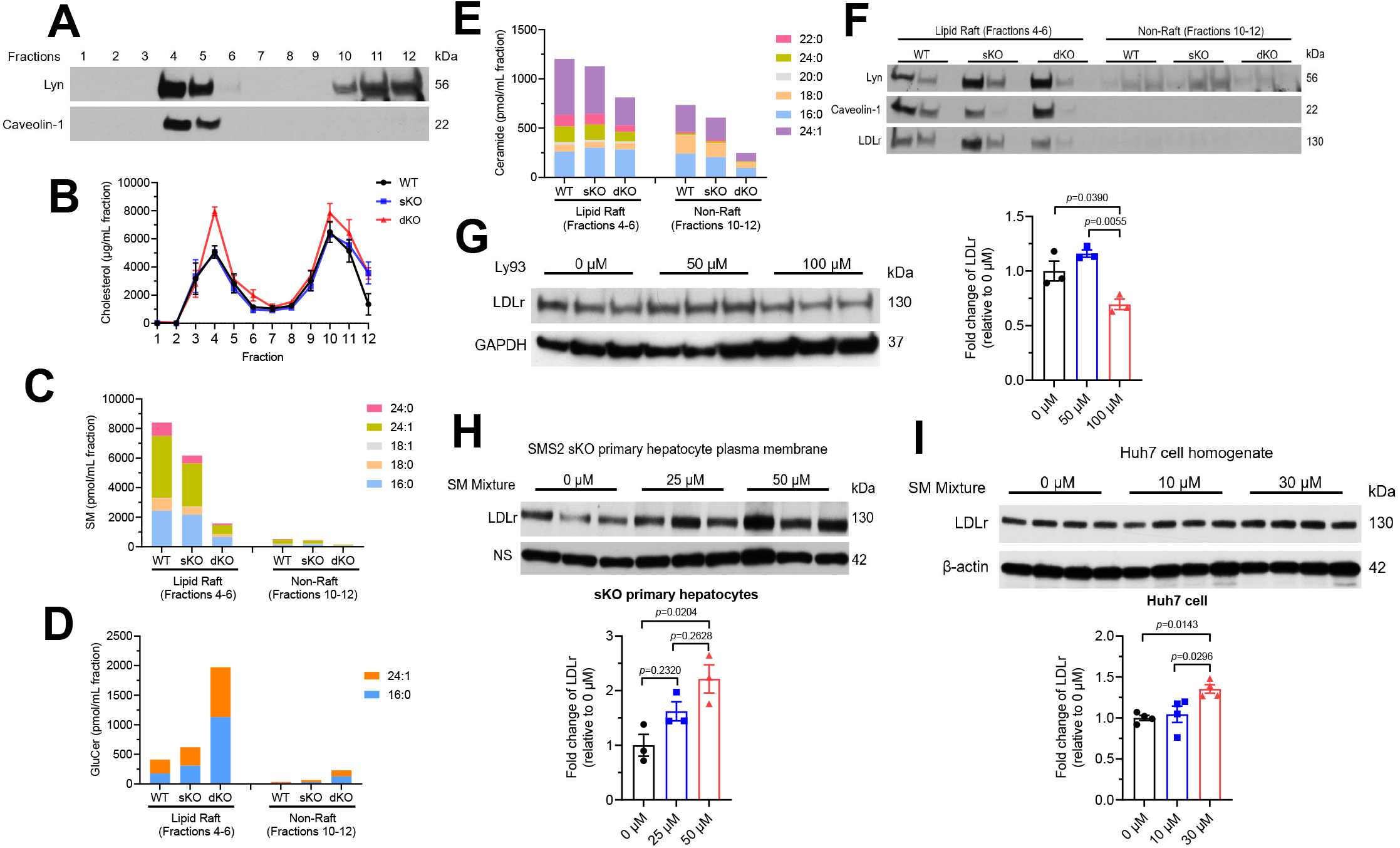
Sphingomyelin reduction decreases LDL receptor expression in hepatocyte lipid rafts. Panel A, Extraction of lipid raft fractions from pooled liver homogenates by sodium carbonate and sucrose density gradient centrifugation (Pooled liver raft fractions from 4 female individual mice per group). Lyn kinase and caveolin-1 were used to detect lipid raft and non-raft fractions. Panel B, Cholesterol levels in fraction 1-12 (Pooled liver homogenates from 4 female individual mice per group). Panels C-E, Sphingomyelin, glucosylceramide, and ceramide levels in lipid raft (Fractions 4-6) and non-raft (Fractions 10-12). Data represent mean±SEM, n=4 individual mice per group. Panel F, Western blot of LDL receptor in lipid raft (Fractions 4-6) and non-raft (Fractions 10-12) from wild-type (WT), *SMS2^−/−^ SMS1^fl/fl^* (sKO), and *SMS2^−/−^ SMS1^fl/fl^ UBC-Cre-ER^T2^* (dKO) mice (Pooled liver raft fractions from 4 female individual mice per group)Panel G, Western blot of LDL receptor in Huh 7 cells supplemented with 0, 50, and 100 μM Ly93 for 18h (n=3 biological replicates per group). Panel H, Western blot of LDL receptor in plasma membrane from female *Sms2* sKO primary hepatocytes supplemented with 0, 25, and 50 μM SM mixture for 28h (n=3 biological replicates per group, primary hepatocytes isolated from male *Sms2* sKO mice). Panel I, Western blot of LDL receptor in Huh 7 cells supplemented with 0, 10, and 30 μM SM mixture for 28h (n=4 biological replicates per group). Panel G through I, Statistical analysis was done using 1-way ANOVA followed by Sidak multiple comparisons test.

Next, we examined LDL receptor in lipid rafts (pooled fractions 4-6 from 4 mice) and non-rafts (pooled fractions 10-12 from 4 mice) and found that the double deficiency reduced LDL receptor in the lipid rafts (**Figure 6F**). We also measured LDL receptor in the fraction #4 and #5 from each mouse liver and found a significant reduction of LDL receptor in all four dKO mouse lipid rafts (**Supplemental Figure 5**). To better understand the molecular mechanism, we treated Huh7 cell, a human hepatoma cell line, with an SM synthase inhibitor (Ly93), and found that 100 mM Ly93 can significantly reduce LDL receptor (**Figure 6G**). We then isolated primary hepatocytes from *Sms2* KO mice and cultured them with or without SM supplementation. We found that SM treatment significantly increases LDL receptor in a dose-dependent manner (**Figure 6H**). Similar results were also obtained from Huh7 cell, a human hepatoma cell line (**Figure 6I**). Interestingly, we treated Huh7 cells with different concentrations of glucosylceramide and ceramide and neither of them could regulate LDL receptor (**Supplemental Figures 6A and 6C**). This was also true for *Sms2* KO primary hepatocytes (**Supplemental Figures 6B and 6D**). Collectively, SM level is one of the determinants of LDL receptor mass in liver cells.

We also examined gene expressions associated with cholesterol metabolism and bile acid metabolism in the liver. As indicated in **Supplemental Figure 7**, we observed that total SMS deficiency led to the significant reduction of *Abcg5, Cyp7a1*, and *Cyp8b1* expressions in a gene dependent fashion mRNA level. However, *Srebp2* and *Sr-b1* expressions were unaffected.

## Discussion

In current study, we prepared inducible global *Sms1* KO/global *Sms2* KO mice and we demonstrated that depletion of *Sms* genes in mice, female or male, resulted in: 1) significant induction of cholesterol levels in the circulation; 2) significant reduction of LDL receptor in the liver; 3) dramatic reduction of SM in membrane lipid rafts, responsible for the reduction of LDL receptor and less LDL be taken up by hepatocytes. Importantly, SM but not ceramide or glucosylceramide treatment induced more LDL receptor expressed in liver cells. This was the very first study indicating the impact of whole-body SM biosynthesis deficiency on cholesterol, especially LDL-cholesterol, metabolism.

One of the key findings of current study was that total SM depletion greatly influenced plasma cholesterol, especially LDL-cholesterol, levels. However, we did not find plasma cholesterol changes in our liver-specific SMS1 KO/global SMS2 KO mice^26^. The potentially reasons could be as follows. First, SMS1 is depleted (mediated by albumin promoter) in early life and this could initial many compensations, while tamoxifen-mediated SMS1 depletion is occurred in adulthood. This is not an uncommon phenomenon. and cell polarity^27,28^. Albumin-Cre-mediated liver-specific deletion of Liver kinase B1 (LKB1) in mice leads to jaundice and lipoprotein-X accumulation in the blood ^29^. However, LKB1 depletion in adult mouse liver had no such effect ^30^. We found that depletion of SPTLC2 (through adenovirus associated virus-Cre) in adult mouse livers did not alter cholesterol or bile metabolism ^31^. However, albumin promoter mediated liver specific SPTLC2 depletion impairs liver function^32^. Second, inducible SMS1 depletion in small intestine and adipose tissue may also contribute to plasma lipid changes as SPTLC2 depletion ^33, 34^.

SM and cholesterol are two important lipids on the monolayer of plasma lipoproteins^15^ and total SMS deficiency caused significant induction of cholesterol in the circulation (**Figure 2E**). This could be due to more production of lipoprotein, especially triglyceride enriched VLDL, from the liver. However, this was not the case. We found that VLDL or triglyceride production was significantly reduced instead of induced (**Figure 2D**) and similar phenomenon was also observed in liver-specific *Sms1* KO/global *Sms2* dKO mice^19^. We also measured plasma triglyceride levels and found that it was reduced (**Figures 2C and 3A**). However, the reduction of VLDL or triglyceride production was not a determinant of cholesterol levels in the circulation. We also found that apoB levels were significantly increased instead of reduced (**Figure 3C**), suggesting that LDL (an apoB-containing and triglyceride depleted particle derived from VLDL) catabolism was altered. Interestingly, we also found increased HDL-cholesterol in the dKO mice (**Figure 3B**) and this was not associated with apoAI (a major apolipoprotein in HDL) levels, since no changes were observed (**Figure 3C**). Similar phenomenon was also observed in liver-specific deficiency of SPT subunit 2, mediated by adenovirus associated virus-Cre recombinase^31^. These results suggested that SM biosynthesis has a specific effect on apoE, particularly on apoE-containing HDL but not apoAI-containing HDL. This phenomenon deservers further investigation. Collectively, total SMS deficiency mediates the formation of a group of triglyceride poor and cholesterol enriched lipoproteins, especially LDL particles. LDL catabolism but not VLDL or triglyceride production determines lipid profile observed in total SMS deficient mice. Then, why those LDL particles can be accumulated in the circulation?

Total SM depletion greatly influences cell membrane lipid composition. Lipid rafts are dynamic microdomains in the plasma membrane. SM and cholesterol are also two partners in lipid rafts, which mediate specific cell functions^35^. Disrupting lipid raft integrity through influencing cholesterol levels perturbs cell-signaling and protein-interaction networks^36^. To investigate the consequence of SM depletion, we separated liver cell membrane into two peaks, lipid rafts and non-lipid rafts. Cholesterol is about evenly distributed in lipid rafts and non-rafts, while SM is mainly located in lipid rafts (**Figure 6B and 6C**). Total SMS deficiency dramatically reduced SM and induced cholesterol levels in the rafts (**Figure 6B and 6C**). SMS uses ceramide to synthesize SM^16^ and we expected that total SMS deficiency should increase ceramide levels in tissues. To our surprise, we found that ceramide levels were reduced instead of induced in the dKO mouse liver cell member lipid rafts and non-rafts (**Figure 6E**), while plasma ceramide levels were increased (**Figure 2B**). Although we could not explain this discrepancy, ceramide levels might have negligible role in mediating LDL catabolism. We have reported that global SMS1 deficiency dramatically induced glucosylceramide accumulation in all tested tissues^17^. SMS1 can use its sterile α-motif to interact with C-terminal of glucosylceramide synthase to form a protein-protein interaction which control tissue glucosylceramide levels^37^. There is no such interaction in SMS1 deficient tissues where glucosylceramide can be synthesized without control. We confirmed this phenomenon (**Figure 6D**). Collectively, total SMS deficiency caused SM depleted, ceramide reduced, glucosylceramide enriched cell, and cholesterol enriched cell membrane.

SM but not glucosylceramide or ceramide levels determine LDL receptor function in liver cell membrane. The LDL receptor pathway for LDL uptake has served as a paradigm for receptor-mediated endocytosis^38^. It has been reported that in HepG2 cells (another human liver hepatoma cell line) the LDL receptor is located in lipid rafts^25^. We found that mouse LDL receptor is located in lipid rafts and SMS-deficiency-mediated SM depletion reduced LDL receptor in them (**Figure 6F, Supplemental Figure 5**). We further found that SM supplementation increased LDL receptor in mouse primary hepatocytes and Huh 7 (**Figures 6H and 6I**). While glucosylceramide or ceramide treatment has no such effect. (**Supplemental Figures 6A-6D**). Collectively, SM is the sphingolipid which responsible for LDL receptor levels and function in the liver.

Although we cannot give a very precise answer for why liver cholesterol is increased (**Figure 4E**) and this induction could contribute to LDL receptor downregulation (**Supplemental Figure 7**), we believe that it should be related with SM but not GluCer or ceramide, since the latter two have no direct effect on cell cholesterol levels ^39,40^. SM and cholesterol are two essential components for lipid rafts formation, and they influence each other ^41–43^. Thus, SMS1 deficiency-mediated SM reduction could be a sensor for more cholesterol production. Thus, increased liver cholesterol could be one of the mechanisms for LDLR downregulation.

Maintaining the balance of cholesterol in circulation depends on the input and output pathways of cholesterol metabolism. The input pathway includes intestinal absorption and the *de novo* synthesis of cholesterol primarily occurred in the liver. The output pathway includes the conversion into bile acids, which are excreted through intestinal tracts and eventually via the feces. Therefore, we also determined pathways other than LDL receptor is(are) influenced by inducible SMS deficiency (**Supplemental Figure 7**). We examined gene expressions which are associated with cholesterol metabolism and bile acid metabolism in the liver. We observed that inducible *Sms1*/global *Sms2* double deficiency led to the significant reduction *Abcg5* mRNA levels, which are associated with cholesterol secretion into bile^44^. Interestingly, we also observed a reduction *Cyp7a1* and *Cyp8b1*, which are the key factors for bile sale production^45^. Collectively, factors other than LDL receptor may also play a role in SMS deficiency-mediated cholesterol metabolism. These phenomena deserver further investigations.

Based on our recent studies on SMS, including current study, we believe that SMS2 but not SMS1 should be a drug target. This is because whole-body *Sms1* KO mice exhibited neonatal lethality and lipodystrophy^46^, while whole-body *Sms2* KO mice are healthy^47^. We and other researchers found that SMS2 deficiency prevented dietary-induced obesity and insulin resistance^47,48^. Moreover, we found that liver SMS2 overexpression promoted fatty acid uptake and liver steatosis, while SMS2 deficiency had an opposite effect compared tp their respective controls^20^. Importantly, we found that SMS2 deficiency attenuated atherosclerosis progression in *Apoe* KO mice^49^.

In summary, inducible whole-body SMS blocking decreases SM in lipid rafts on liver cell membrane and decreases LDL receptor mass and LDL uptake, thus, increasing cholesterol in the circulation.

## Abbreviations

SMS: sphingomyelin synthase
LDL: low-density lipoprotein
KO: gene knock out
dKO: SMS1/SMS2 double KO
WT: wild type

## Acknowledgments

This work was supported by VA Merit 000900-01, NIH HL139582, and NIH RO1HL149730 grants to Xian-Cheng Jiang. We thank Dr. Deyong Ye for providing Ly93.

## Author Contributions

Z.L. and M.H. performed 90% of the experiments, analyzed data, and modified the manuscript. G.C. performed some real-time PCR and western blots. T.S.W. measured sphingolipids using LC/MS/MS. X.C.J. conceived the ideas, designed, and discussed experiments, supervised progress, and wrote the manuscript.

## Significance

SM and cholesterol are two key lipid partners on cellular membranes where they form specific microdomains, called lipid rafts. Lipid rafts provide a platform for mediating specific cell functions, including lipoprotein up-taking, such as low-density lipoprotein (LDL) receptor mediated LDL catabolism. SM and cholesterol are also two key lipids on the monolayer of plasma lipoproteins, including LDL, which participate in lipid transport in the circulation. Many studies have indicated the influence of cholesterol on SM, however, the influence of SM on cholesterol is still not well known. The current study examined the influence of SM biosynthesis on the metabolism of cholesterol, especially LDL-cholesterol. The results obtained not only related to the biology of SM but also related to future therapeutic approach designing, in terms the treatment of metabolic diseases.

